# Simulation study evaluating the ability of two statistical approaches to identify variance quantitative trait loci Arabidopsis and maize

**DOI:** 10.1101/2021.06.25.449982

**Authors:** Matthew D. Murphy, Samuel B. Fernandes, Gota Morota, Alexander E. Lipka

## Abstract

Genomic loci that control the variance of agronomically important traits are increasingly important due to the profusion of unpredictable environments arising from climate change. The ability to identify such variance quantitative trait loci (vQTL) in association studies will be critical for future breeding efforts. Two statistical approaches that have already been used to detect vQTL are the Brown-Forsythe test (BFT) and the double generalized linear model (DGLM). To ensure that they are deployed to variance genome-wide association studies as effectively as possible, it is critical to study the factors that influence their ability to identify vQTL. We used genome-wide marker data in maize (*Zea mays* L.) and *Arabidopsis thaliana* to simulate traits controlled by variance quantitative trait nucleotides (vQTNs) and then quantified true and false positive detection rates of the BFT and DGLM. We observed that the DGLM yielded similar or higher true positive vQTN detection rates than the BFT, regardless of the effect size or minor allele frequency (MAF) of the vQTNs. Low true positive detection rates were noted for QTNs with low MAFs (~0.10), especially when tested on subsets of *n* = 500 individuals. We recommend that larger data sets than those used in our study (i.e., *n* > 2,532) be considered to overcome these low observed true positive detection rates. Such an undertaking should maximize the potential of the BFT and DGLM to highlight which vQTLs should be considered for further study.

## Introduction

The world’s food baskets face an expanding amount of unpredictable growing seasons due to the ongoing threat of climate change (Ziervogel and Erickson, 2010). If a 4°C increase in global temperature is not prevented by 2100, there could be a potential loss of $23 quadrillion to agriculture (Schillaci *et al.*, 2019). Unfortunately, most crops are maladapted to highly variable environments, where optimal growing conditions may never be attained (Mulder *et al.*, 2007). An idea that may lend itself to accelerating the development of crops better suited for such variable environments is canalization. Canalization is the hypothesis that natural selection minimizes variation for certain traits in a way that prevents major loci from being influenced significantly by the environment or background genetic variance like epistasis (Ronnegard and Valdar, 2011). Artificial selection facilitates the decanalization of certain loci, which has allowed domesticated crops to grow in novel environments (Kitano, 2004). However, these decanalized loci are disadvantageous if the environment it was adapted to becomes unpredictable (Waddington, 1942). Collectively, the combination of decanalized loci and unpredictable environments has resulted in such loci controlling the variance of a targeted trait (Debat & David, 2001). A classic example of such variance-controlling loci are genes that encode heat shock proteins, which are involved with various environmental stressors, including heat stress, ultraviolet radiation, cold tolerance, and biotic stressors (Park and Seo, 2015). Variance-controlling loci also arise from epistatic gene action, where the marginal effects of one of the epistatically interacting genes appear as a vQTL (see Forsberg and Carlborg 2017 for a review). While current genome-wide association study (GWAS) approaches can indirectly identify these variance-controlling loci through the inclusion of additional gene by environment (GxE) and epistatic (GxG) terms in the model, they are not able to directly quantify their effects (Shen *et al.*, 2014; Yadav *et al.*, 2016). Thus, variance GWAS (vGWAS) has been proposed to quantify the effect these variance-controlling loci (Shen *et al.*, 2012).

The primary purpose of vGWAS is to detect genetic loci that alter the variance of a phenotype between different genotypes (Al Kawan *et al.*, 2018). While vGWAS has been conducted in plants and crops, its utilization is still not widespread. To date, vGWAS has been conducted for ionomic traits, including molybdenum content in *Arabidopsis thaliana* and cadmium content in bread wheat *(Triticum aestivum)* (Shen *et al.*, 2012; Forsberg *et al.*, 2015; Hussain *et al.*, 2020), as well as for oil-related traits in maize (*Zea mays* L.) (Li *et al.*, 2020). Unlike those used in a standard GWAS (denoted as a mean GWAS or mGWAS), the statistical models used for vGWAS specifically assume unequal phenotypic variance at each genotypic state of a given locus, i.e., in the presence of variance heterogeneity (Ronnegard and Valdar, 2011). Variance-controlling loci are connected to many different ideas within quantitative genetics, including epistatic and GxE interactions (Struchalin *et al.*, 2012; Rönnegård and Valdar, 2011). The advantages of using vGWAS for such interactions include i) bypassing the interaction term often used in the statistical models for these interactions and ii) prioritizing genomic regions likely to harbor epistatic interactions, thereby reducing the severity of multiple testing correction (Struchalin *et al.*, 2012; Petterson and Carlborg, 2015).

Many statistical analyses have been developed to test for variance heterogeneity. From a biological perspective, the choice of test and model can be divided into whether or not one accounts for population structure, relatedness, and other covariates (Ronnegard *et al.*, 2012). Of the statistical tests that do not account for such factors, Levene’s test and its median modification, the Brown-Forsythe test (BFT), have been the most popular (Brown and Forsythe, 1974; Rönnegård and Valdar, 2012). Although they are useful as a quick diagnostic for identifying variance-controlling loci, they cannot explicitly correct for population structure, familial relatedness, or for non-variance controlling loci (called mean QTLs or mQTLs) (Hong *et al.*, 2017). In contrast, models that allow for the inclusion of these factors as covariates theoretically offer higher power to detect variance-controlling loci. In particular, the double generalized linear model (DGLM) (Lee and Nelder, 1996) adjusts for potential confounding between vQTLs, mQTLs, and population structure through the inclusion of fixed-effect covariates. Excitingly, more sophisticated versions of the DGLM also include random effects to account for confounding due to familial relatedness (Lee and Nelder, 2006; Rönnegård and Valdar, 2012).

Although statistical approaches seeking to estimate the effects of variance-controlling loci have opened up many opportunities for discovering new sources of quantitative trait variation, detecting variance-controlling loci still poses challenges. For example, the precision needed to detect a variance-controlling locus often requires five times as many individuals compared to the precision needed to detect a mean-controlling locus (Lee and Nelder, 2006; Ronnegard and Valder, 2012). This suggests that there is a critical need to systematically study the statistical performance of leading vGWAS approaches. Therefore, the purpose of this study was to explore the factors that influence the ability of the BFT and DGLM to detect vQTLs underlying plant traits. We used publicly available genotyping-by-sequencing (GBS) marker sets from the 1,001 genomes diversity panel in *Arabidopsis thaliana* (Alonso-Blanco *et al.*, 2016) and the USDA-ARS North Central Region Plant Introduction Station (NCRPIS) Panel in *Zea mays* L. (Romay *et al.*, 2013) to simulate traits controlled by variance-controlling loci with various effect sizes. We hypothesized that that accounting for mQTLs in the statistical approach would increase the ability to detect vQTLs. Accordingly, we predicted that whenever mQTLs and vQTLs both contributed to the genetic signal of a simulated trait, the DGLM would yield higher true positive vQTN detection rates than the BFT.

## Materials and Methods

### Genotypic data and filtering procedures

We conducted simulation studies using genotypic data from two plant species with contrasting levels of linkage disequilibrium (LD) decay. The first genotypic data set was a subset of 1,087 accessions from the *Arabidopsis thaliana* 1,001 genomes diversity panel, available at https://1001genomes.org/ (Alonso-Blanco *et al.*, 2016). The 1,001 genomes diversity panel consists of germplasm mostly collected from Eurasia, North America, and Northern Africa. These accessions were genotyped, using GBS, which produced 10,707,430 biallelic SNPs (Alonso-Blanco *et al.*, 2016). The second set of genotypic data was a subset of 2,532 lines from the NCRPIS diversity panel in maize (Romay *et al.*, 2013). This diversity panel was genotyped for 681,257 SNPs, as described in Romay *et al.* (2013). These genotypic data are publicly available at cbsusrv04.tc.cornell.edu/users/panzea/download.aspx?filegroupid=6.

Both genotypic data sets were filtered by removing SNPs with more than 10% and 5% missing data in *Arabidopsis* and maize, respectively, using VCFtools (Danecek *et al.*, 2011). We also used VCFTools filter by minor allele frequency (MAF) of 5% and 1% in *Arabidopsis* and maize, respectively. These data sets were then further filtered with LD pruning utilizing PLINK (Purcell *et al.*, 2007). A different criterion for LD pruning was used to obtain a similar number of SNPs between the two species. In *Arabidopsis*, the LD pruning parameters for the 1,001 Genomes GBS data set was r^2^ = 0.10, a window size of 200 kb, and a step size of 20 kb. For maize, these parameters were set to r^2^ = 0.90, with a window size of 100 kb and a step size of 10 kb; similar parameter values were used in a previous simulation study (Fernandes *et al.*, 2021). The resulting number of remaining SNPs left for *Arabidopsis* and maize were 41,384 and 41,611, respectively.

To assess how sample size affects the performance of the tested statistical methodologies, we considered two different sample size scenarios for each species. The first scenario focused on employing all individuals in both panels (i.e., 1,087 for *Arabidopsis* and 2,532 for maize). In the second scenario, we randomly selected *n* = 500 individuals from each panel using the sample() function in R (R Core Team 2021).

### Simulating traits controlled by vQTNs

For a given setting (i.e., configuration of mean QTNs, variance QTNs, their effect sizes, and narrow-sense heritability), the simulated trait values for each individual were calculated from the following formula (Hill and Mulder, 2010):

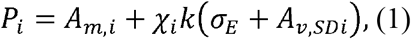

where *P_i_* is the simulated phenotypic value of the *i^th^* individual, *A_m,i_* is the collective genetic value from all simulated mean QTNs for the *i^th^* individual *χ_i_*, is a standard normal random variable (i.e., *N*(*μ* = 0, *σ*^2^ = 1)) sampled for the *i^th^* individual, *k* is a constant described in the paragraph below to enable a certain degree of control over the narrow-sense heritability, *σ_E_* is the population standard deviation determined attributed to non-genetic sources, and *A_v,SDi_* is the collective genetic value of all simulated vQTN for the *i^th^* individual. The values of *A_m,i_* and *A_v,SDi_* are respectively calculated as the sum of observed genotype codes at each mean and variance QTN, multiplied by the (respective) mean and variance QTN effects for the *i^th^* individual. These simulations are conducted assuming that the covariance between *A_m,i_* and *A_v,SDi_* is zero.

One major challenge for simulating traits controlled by vQTNs is the specification of the desired heritability. Because the value of (*σ_E_* + *A_v,SDi_*) changes for every individual, the value of the heritability will also change for every individual. We, therefore, made *ad hoc* adjustments to (1) to ensure at least partial control for a desired narrow-sense heritability (*h*^2^). First, the value of *σ_E_* was also set to 1, and then *A_m,i_* was centered and scaled, so its sample mean and standard deviation were respectively 0 and 1. These steps were taken to facilitate the estimation of the *k* in (1).

We now describe the derivation of the procedure we used to estimate the value of *k.* Consider the following modified formula for estimating narrow-sense heritability *h*^2^ for traits controlled by vQTNs:

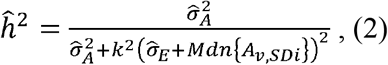

where 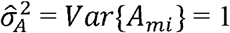 because *A_m,i_* was scaled, 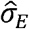 was set equal to *σ_E_* = 1 to facilitate calculations, and *Mdn*{*A_v,SDi_*} is the median value of *A_v,SDi_* across all *n* individuals (i.e., all individuals in either the *Arabidopsis* or maize data sets used for the simulations). Thus, solving (3) for *k* yields:

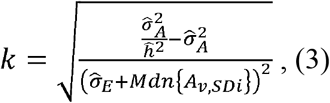

where all terms are as previously described. Thus, for each simulation setting, the value of *k* estimated from (3) was used in (1) to obtain simulated trait values for every individual.

Two R functions were used to simulate these traits. Additive mean QTNs, which contribute to *A_m,i_* in (1), were simulated using the create_phenotypes() function in the simplePHENOTYPES R package (Fernandes and Lipka, 2020). We then developed our custom R function that was roughly based on the Python code from Dumitrascu *et al.* (2018) to obtain the remaining necessary values in (1), (2), and (3) to simulate the phenotypic values *P_i_*. To facilitate the deployment of our simulation pipeline to future studies, we made it available through simplePHENOTYPES v1.4 (create plienotvpes(…, model = “V”)) (https://github.com/samuelbfernandes/simplePHENOTYPES).

### Specific genetic architectures considered in five settings

We simulated traits under five different settings within each species and sample size scenarios. These settings are based on previous vGWAS and vQTL studies conducted in *Arabidopsis* and maize (Shen *et al.*, 2012; Li *et al.*, 2013; Forsberg *et al.*, 2015; Li *et al.*, 2020). The specific number of additive and variance QTNs, their effect sizes, and the overall narrow-sense heritability (*h*^2^) varied across these settings and are summarized in Table 1. Briefly, to enable a rigorous assessment of false-positive rates of the tested vGWAS approaches, the first setting (“Null” on Table 1) consisted of traits with a heritability of *h*^2^= 0 and zero QTNs. The purpose of the next two settings (“Oil-like MAF 0.10” and “Oil-like MAF 0.40”) was to simulate two genetic architectures that were identical in all aspects except for the targeted minor allele frequency of the vQTNs. The corresponding genetic architectures were derived from what is currently known about oil content in maize, a trait of great economic importance and well-studied within quantitative genetics (Li *et al.*, 2020). The last two settings (“Mo-like” and “Mo-like without mQTN”) were created to explore the impact of the presence of mQTNs on true positive detection rates. The genetic architecture of these two was set to resemble the findings of Shen *et al.* (2012) when they searched for vQTL associated with molybdenum content in *Arabidopsis.* Within each species, a total of 100 replicate traits were simulated at each setting.

**Table 1:** Description of the input parameter values for each trait in our simulation studies.

| Simulation Setting | $h^2$ | Targeted<br>MAF of<br>QTNs | # Mean QTN | Mean QTN<br>effect size | # Variance<br>QTN | Variance QTN<br>effect size |
| --- | --- | --- | --- | --- | --- | --- |
| "Null" | 0 | NA | 0 | NA | 0 | NA |
| "Oil-like MAF 0.10" | 0 | 0.10 | 0 | NA | 17 | 0.10-0.90<sup>a</sup> |
| "Oil-like MAF 0.40" | 0 | 0.40 | 0 | NA | 17 | 0.10-0.90<sup>a</sup> |
| "Mo-like" | 0.63 | 0.40 | 1 | 0.25 | 1 | 0.90 |
| "Mo-like without mQTN" | 0 | 0.40 | 0 | NA | 1 | 0.90 |

$h^2$ narrow-sense heritability, *MAF* minor allele frequency, *QTN* quantitative trait nucleotide, *NA*
not applicable
<sup>a</sup> The effect sizes of these 17 vQTNs increased between 0.10 and 0.90 in increments of (0.90-0.10)/17 ≈ 0.0471.

### Competing vGWAS models and tests

We considered two different statistical approaches used in previous plant publications to conduct vGWAS, namely the BFT and the DGLM (Shen *et al.*, 2012; Forsberg *et al.*, 2015; Hussain *et al.*, 2020; Li *et al.*, 2020). The BFT is used to test for variance homogeneity (Brown and Forsythe, 1974; Shen *et al.*, 2012). For each locus, the BFT evaluates:

*H*_0_: *Population variances of traits are equal at all genotypes* versus
*H_a_: Population variances of traits are different for at least one genotype*,

and uses the corresponding test statistic:

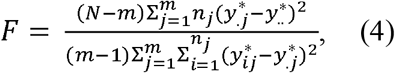

where *N* is the total number of accessions, *n_j_*, is the number of accessions in the genotypic group *n_j_, m* is the number of allelic states of a genetic marker, and

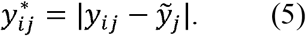

In (5), *y_ij_* is the phenotypic value for the *i^th^* accession with the *j^th^* genotype and 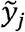 is the median phenotypic value of individuals with genotype *j*. Under *H*_0_, the BFT statistic in (4) follows an *F* distribution with degrees of freedom equal to *m* – 1, *n* – *m* (Shen *et al.*, 2012). The BFT was performed using the brown.forsythe.test function from the vGWAS R package (Shen *et al.*, 2012). Because the BFT does not allow explicit inclusion of covariates to account for false positives arising from population structure and familial relatedness, it often serves as a quick diagnostic test to see if the trait of interest has any underlying vQTLs. Furthermore, the BFT is robust to phenotypic departures from normality (Dumitrascu *et al.*, 2019; Hussain *et al.*, 2020).

The DGLM belongs to a family of generalized linear models, which relaxes the assumption of normality of phenotypic residuals for more flexible modeling. Specifically, the DGLM is subdivided into two components that model the relationship between a response variable and i) explanatory variables controlling its population mean, and ii) explanatory variables controlling its population variance. The component of the DGLM controlling the population mean is written as follows:

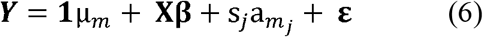

where ***Y*** is an *n*-dimensional vector representing the phenotypic response variable, μ_*m*_ is the intercept; **1** is an *n*-dimensional vector of 1’s, ***X*** is an *n* x *q* design matrix containing the first *q* principal components from a principal component analysis (PCA) of the markers (which was 4 and 3 for *Arabidopsis* and maize, respectively) (Price *et al.*, 2006); ***β*** is a *q*-dimensional vector of the regression coefficient for the principal components; *s_j_* is an *n*-dimensional vector containing the *j^th^* SNP encoded as 0,1,2 at the *j^th^* marker, *a_m_j__* is the effect size of the *j^th^* marker, and 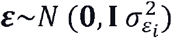 is an *n*-dimensional vector of normally distributed error terms. In the simulation setting called “Mo-like” in Table 1, *s_j_* is set equal to the mQTN and *a_m_j__*. is its effect size; in all other settings, these two terms are omitted from the model because no mQTNs were simulated. The component of the DGLM controlling the population variance of the *i^th^* individual 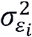 is written as follows:

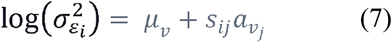

where 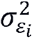 is the residual variance for genotype *i*; *μ_v_* is the intercept, *s_ij_* is the observed SNP value encoded 0,1 and 2 at the *j^th^* marker for genotype *i*, and *a_v_j__* is the effect size of the *j^th^* marker. To test for a significant association between the *j^th^* marker and the variance of the tested trait, we used the Wald test (Agresti 2003) to test *H*_0_: *a_v_j__*. = 0 versus *H_a_*: *a_v_j__* ≠ 0, which follows an asymptotic 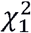 distribution under *H*_0_. To perform DGLM, we used the dglm R code from Hussain *et al.* (2020), which came from the dglm R package (Dunn and Symth, 2020). The PCAs for population structure were obtained using GAPIT version 4.0 (Lipka *et al.*, 2012).

The ensuing analyses using both of these statistical approaches were conducted on a Dell Precision Tower 3240 with 64.0 GB RAM. While the BFT was ran on a single core, DGLM was ran on four cores using the foreach R package.

### QTN detection rates for competing models

To assess whether or not the vGWAS methodologies can correctly identify markers as associated with our simulated traits, we evaluated the true and false-positive QTN detection rates using the Benjamini-Hochberg (1995) procedure to control the false-discovery rate (FDR) at 5%. A statistically significant SNP was labelled as a true positive if it was within a 250k window of a simulated vQTN for maize (Lipka *et al.*, 2013) and within 100k window of a simulated vQTN in *Arabidopsis* (Gonzalez-Jorge *et al.*, 2013). Likewise, a statistically significant SNP was labelled as a false positive if it was outside of these windows. We measured the true-positive rate as the proportion of times we detected at least one true-positive per replication out of 100 replications. Similarly, our false-positive rate is defined as the proportion of times we detected at least one false-positive per replication out of 100 replications.

## Results

### False positive detection rates in the “Null” setting suggest BFT and DGLM adequately control for false positives

We ran the “null” settings in both species to verify that the observed false-positive rates for the BFT and DGLM were similar to what we would expect based on statistical theory. Across both species and sample sizes considered, the observed false-positive rates for both approaches were reasonably close to the theoretical value of 0.05 (Fig. 1). This suggests that both the BFT and DGLM were adequately controlling for false positives.

**Fig. 1.**
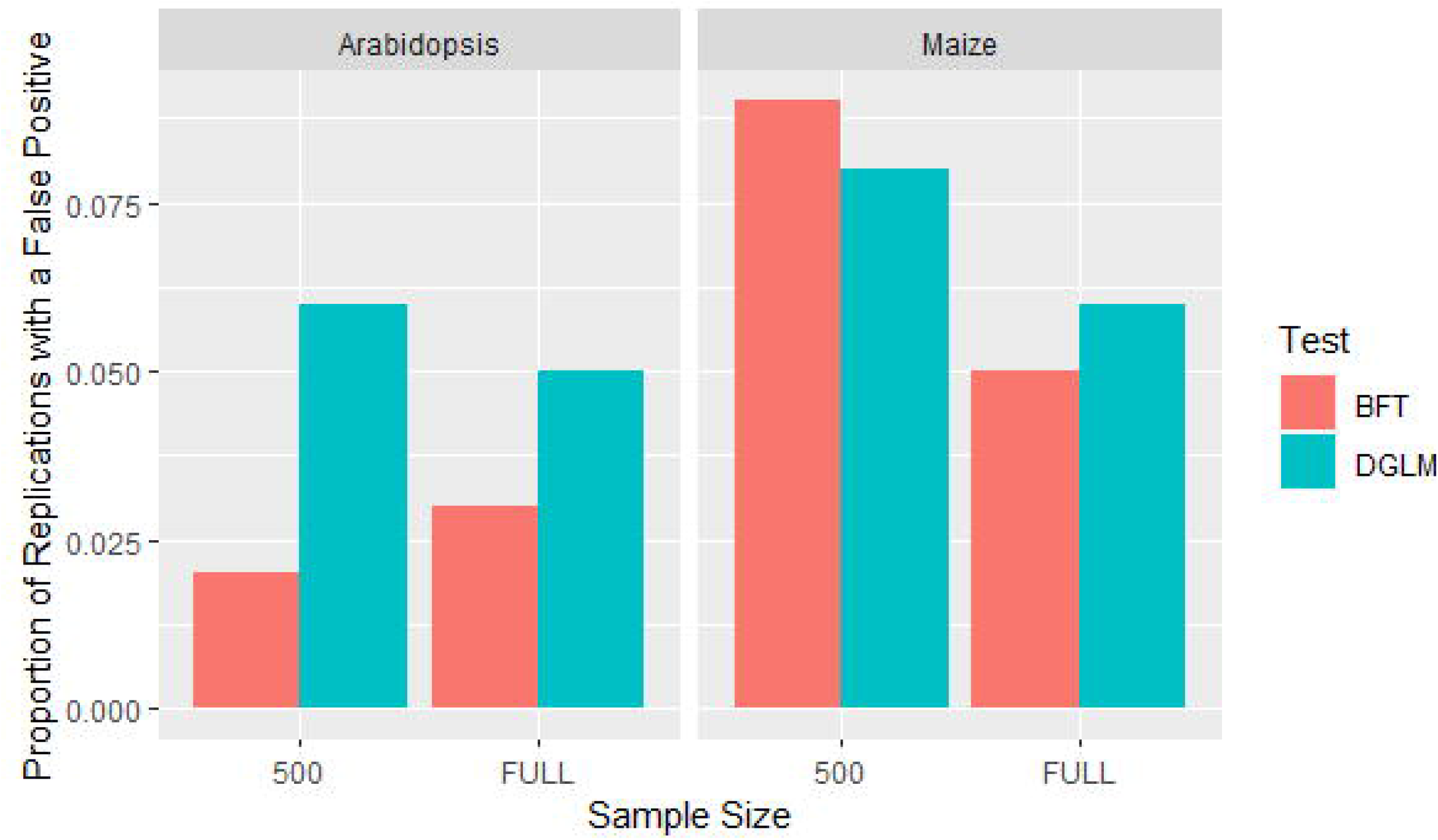
False-positive detection rates for the null setting at a false discovery rate of 0.05. The x-axis represents the sample size of each diversity panel. The y-axis is the proportion of replications where a false positive is detected at least once. Each panel represents the species. BFT: Brown-Forsythe test; DGLM: double generalized linear model

### DGLM yielded similar or higher true positive detection rate than BFT, regardless of the presence of mQTNs

To test the hypothesis that accounting for mQTNs would increase the ability to detect vQTNs, we compared the observed true positive vQTN detection rates of the BFT and DGLM in the “Mo-like” and “Mo-like without mQTN” settings. If this hypothesis were true, we would have expected to see i) the DGLM yielding higher true positive detection rates than the BFT in the “Mo-like” setting, and ii) the discrepancy in true positive detection rates between DGLM and BFT to be more pronounced in the “Mo-like” setting. For both species, we observed that the DGLM yielded true positive detection rates that were greater than or equal to those of the BFT at both settings, which supports the prediction in i). However, the discrepancy in the true positive detection rates was not more pronounced in the “Mo-like” setting (Fig. 2). Although this result does not support our hypothesis, it does suggest that the DGLM is capable of yielding higher detection rates than the BFT, regardless of whether or not mQTNs also underlie the genetic signal of the simulated trait.

**Fig. 2.**
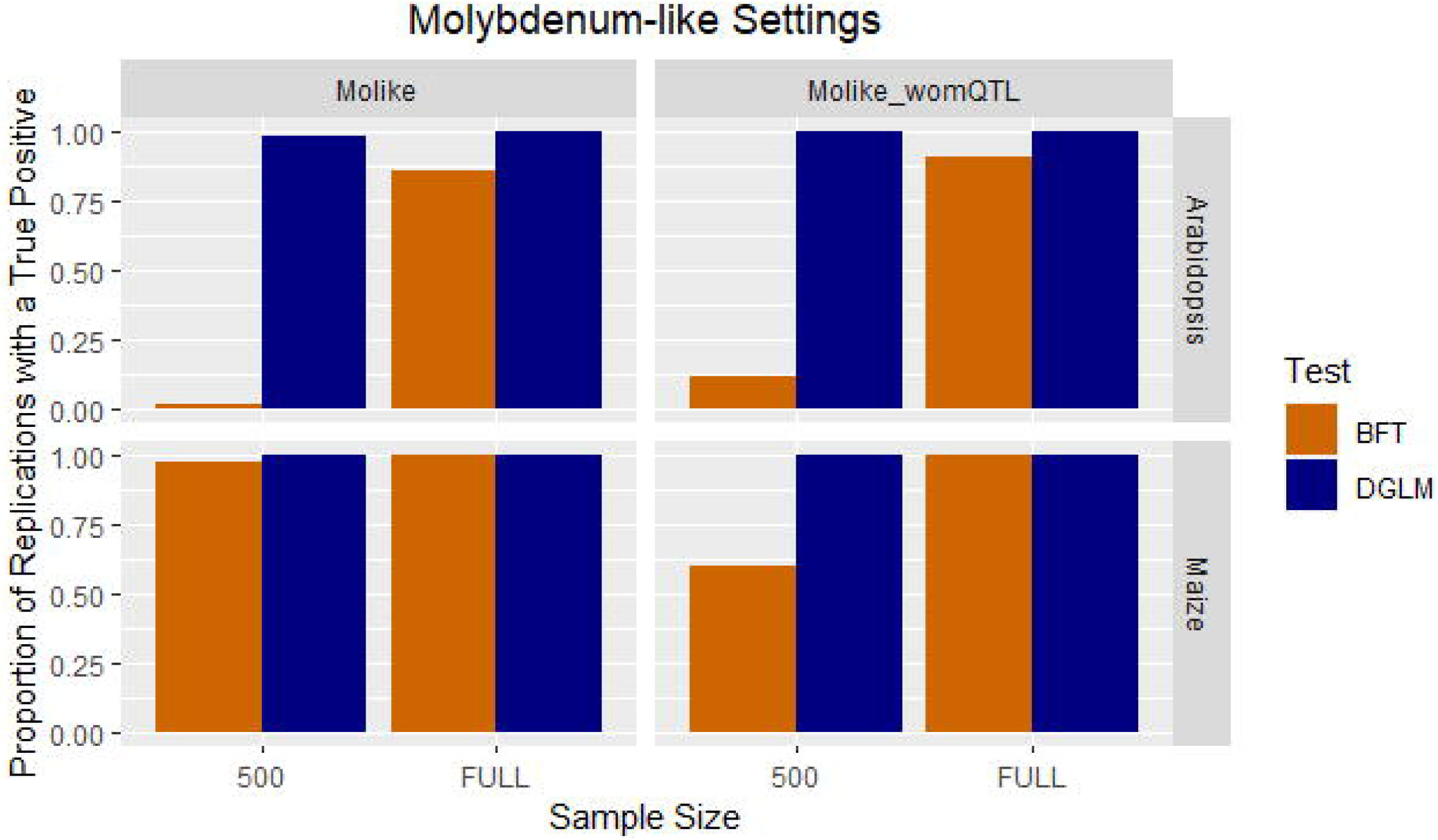
True-positive detection rates for the molybdenum-like settings at a false discovery rate of 0.05. The x-axis represents the sample size of each diversity panel. The y-axis is the proportion of replications where a true positive is detected at least once. The upper and lower panels represent *Arabidopsis* and maize, respectively. The left and right panels represents the variation of the Molybdenum-like setting. BFT: Brown-Forsythe test; DGLM: double generalized linear model

### True positive detection rates were sensitive to MAFs of vQTN but not to their effect sizes

The two “oil-like” settings were conducted to enable an evaluation of true positive detection rates of the BFT and DGLM across vQTNs that differ in effect sizes and MAFs. Within each species and sample size considered, the true positive detection rates of the BFT and DGLM were relatively consistent across the simulated effect sizes, with DGLM consistently yielding higher true positive detection rates than the BFT (Fig. 3). In contrast, we noted substantial differences in true positive detection rates across the different targeted MAFs that were considered. Notably, substantially lower true positive detection rates were observed whenever the targeted MAFs of the selected vQTNs were 0.10. This result suggests that the ability of both the BFT and DGLM are highly dependent on the MAF of the simulated vQTNs.

**Fig. 3.**
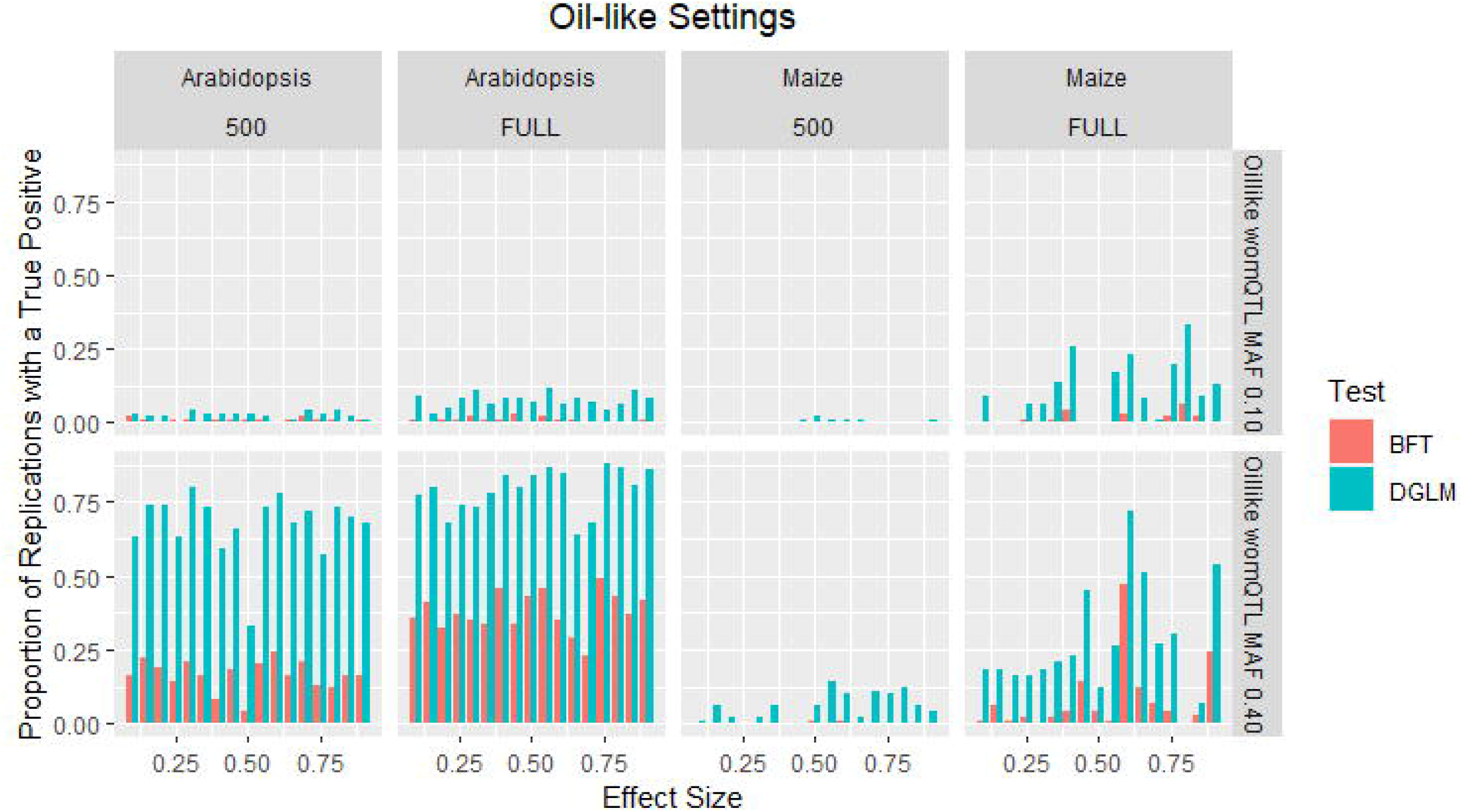
True-positive detection rates for the oil-like settings at a false discovery rate of 0.05. The x-axis represents the effect sizes of each species and sample size combination. The y-axis is the proportion of replications where a true positive is detected at least once. The upper and lower panels represent the oil-like setting. Each column represents the species and trait combination. BFT: Brown-Forsythe test; DGLM: double generalized linear model

### Larger sample sizes are needed to detect vQTNs with low MAF

Evaluation of the performance of the BFT and DGLM at the smaller sample size of *n* = 500 was critical for determining the extent to which these approaches could facilitate the elucidation of variance-controlling loci when researchers are constrained with small data sets. Unsurprisingly, lower true positive detection rates were observed at *n =* 500 in both species, with the BFT yielding substantially smaller true positive detection rates than the DGLM.

At the setting where the targeted MAF of the vQTNs were 0.10 (i.e., “Oil-like MAF 0.10”), we consistently observed low true positive detections across both species at *n* = 500, with only marginal increases in detection rates at the full sample sizes of *n* = 1,087 *Arabidopsis* accessions and *n* = 2,532 maize lines (Fig. 3). Collectively, these results suggest that substantially larger sample sizes than those considered in our study are needed for the BFT and DGLM to detect vQTNs with low MAFs with reasonable statistical power.

## Discussion

We used genome-wide marker data in maize and Arabidopsis to simulate traits controlled by vQTNs, and then compared the resulting true and false positive vQTN detection rates from the BFT and the DGLM. While both of these vGWAS approaches adequately controlled the false positive detection rates, we found that the DGLM yielded true positive detection rates that were greater than or similar to those from the BFT, regardless of the presence of mQTNs. We also observed that across all of the sample sizes we considered, true positive detection rates were low for vQTNs with MAFs that were approximately 0.10. These results provide a potential benchmark for how the BFT and DGLM are expected to perform when deployed to vGWAS in plants. Such an assessment is essential because the more widespread use of vGWAS in plants could substantially facilitate breeding for uniformity of trait values across various environmental conditions.

### Implications of simulation results on the use of BFT and DGLM in plant vGWAS

One of the most surprising results was the relative stability of true positive vQTN detection rates across the various effect sizes that were simulated. Given that this finding was somewhat counterintuitive, we recommend that future studies explore the impact of vQTN effect size across more species, sample sizes, and genetic architectures.

Nevertheless, our results underscored the general finding that the ability of current vGWAS methodologies to identify vQTNs is not as high as their additive counterparts (as shown in, e.g., Rönnegård and Valdar, 2011). In particular, we showed the extent to which true positive vQTN detections could decrease for the BFT and DGLM when sample sizes are small. Importantly, our simulations clearly demonstrated that a sample size of *n* = 500 is inadequate for both the BFT and the DGLM to identify vQTNs with MAFs that are at most 0.10. Regardless of diversity panel size and species, we also observed that true positive vQTN detection rates were consistently higher at MAFs of approximately 0.40. Given that diversity panels are very likely to contain rare causal variants (Lipka *et al.*, 2015), it will be critical to take measures to ensure that the influence of MAF on true positive vQTN detection rates is mitigated in practice. The first measure we advise taking is to maximize the sample size. Second, we recommend deploying vGWAS to panels and data sets where the MAFs of causal mutations are substantially higher than 0.10, as we would expect both the BFT and DGLM to yield substantially higher vQTN detection rates.

Another surprising result was that we did not observe clear evidence that accounting for mQTNs increased the ability to identify vQTNs. Instead, we observed that DGLM (which accounts for the effects of mQTNs) uniformly yielded high true positive detection rates in two nearly identical simulation settings that only differed in whether or not an mQTN was present (i.e., the “Mo-like” and “Mo-like without mQTN” settings described in Table 1). Within each species, we similarly observed stable true positive vQTN detection rates from the BFT (which does not account for the effects of the mQTNs) across these two simulation settings. These results align with those observed in a similar study in biparental mice populations (Corty and Valdar 2018). However, it is important to note that our mQTN had a relatively small effect size of 0.25. As the variance genetic architectures of more traits are elucidated, future simulation studies should focus on mQTNs with larger effect sizes and multiple mQTNs instead of our single mQTN with a small effect size.

### Impact of computational time on running the BFT and DGLM

Depending on the sample size, we observed that the time required for the DGLM to complete a vGWAS of approximately 40,000 markers ranged from approximately 9 (for *n* = 500 *Arabidopsis* and maize individuals) to 32 minutes (for *n* = 2,532 maize individuals). In contrast, we noted that the BFT completed a vGWAS on the same data sets in approximately three to six minutes. Thus, the increase in computational time required to complete the DGLM relative to the BFT was relatively modest for our genotypic data sets and sample sizes. We argue that for data sets of a similar scale, the extra computational time of the DGLM is worthwhile because we would expect this approach to yield higher true positive detection rates compared to the BFT.

For larger data sets, we recommend implementing parallelization when performing DGLM. However, if the computational time required for completing the DGLM still makes its usage infeasible even after parallelization, then a two-step procedure similar to that of Cordeva-Polmera et al. (2020) utilizing both statistical approaches could be conducted. That is, a genome-wide scan using the BFT could first be undertaken to identify peak-associated markers, and then the DGLM could be conducted on only such markers identified by the BFT.

### Limitations of our simulation studies

Although useful for simulating traits with similar genetic architectures of real traits, the approach we implemented to account for the narrow-sense heritability was *ad hoc.* We recommend that future studies focus on accounting for broad-sense heritabilities, as this would enable more user-control over the total phenotypic variance attributable to genetic effects. Another drawback is that our study did not consider investigating the relationship between vQTNs and epistasis. Given that previous simulation studies already investigated this relationship (Dumitrascu et al., 2019; Forsberg and Carlborg, 2017), further exploration would reveal the extent to which a preliminary vGWAS in plants could pinpoint genomic regions most likely to harbor epistatically interacting loci.

Another limitation of our study is that we explored only one configuration of modeling the relationship between vQTNs and a trait. Specifically, the configuration we used in (1) is based on the standard deviation additive model (Hill and Zhang, 2004; Hill and Mulder, 2010). Other vQTN quantitative genetics models, such as reaction norm model from Hill and Mulder (2010), could be used to simulate more scenarios where vQTNs could arise including phenotypic plasticity, GxE interactions, and epistasis.

### Areas for future research

The inbreeding species considered in our simulation studies, *Arabidopsis*, has not been subjected to as much artificial selection compared to what would be expected in crops (Izawa, 2007; Woodward and Bartel, 2018). Thus, future simulations should consider an inbreeding crop species such as sorghum (*Sorghum bicolor* L. Moench) or rice (*Oryza sativa* L). Additionally, our decision to simulate traits similar to those where vQTN have already been identified resulted in our simulated traits resembling metabolic traits with tractable genetic architectures. However, a recent report in maize showed that vQTNs are also present in plant architectural and phenology traits (Zhang and Qi, 2021). Thus, the practicality and utility of the BFT and DGLM to identify vQTNs in crops could be more comprehensively explored if a wider range of genetic architectures were studied in future work.

While the BFT and DGLM are two commonly used statistical methodologies for testing for vQTNs, there are other statistical approaches that could be incorporated into vGWAS. Two such approaches are the hierarchical generalized linear model (HGLM) (Lee and Nelder, 1996) and double hierarchical generalized linear model (DHGLM) (Lee and Nelder, 2006). These models account for familial relatedness by including the individuals as a random effect and setting their variance-covariance to be proportional to an additive genetic relatedness matrix. Although the associated computational complexity of fitting these two models rendered them impractical to evaluate in our simulation studies, they have been previously evaluated in wheat (Hussain et al. 2020) and animal breeding (Rönnegård et al., 2010). Given that the DGLM and HGLM in Hussain et al. (2020) both identified the same vQTNs associated with cadmium content in wheat, we would expect that the HGLM and DHGLM to yield similar true positive vQTN detection rates for traits that are not associated with familial relatedness. We recommend that future work focuses on increasing the computational efficiency of the HGLM and DHGLM so that their ability to identify vQTNs could be studied in a manner similar to that which is presented in this work.

## Conclusion

The ability of vGWAS approaches to identify vQTN needs to be thoroughly scrutinized before they can become more commonplace in quantitative genetics analysis in plants. Based on our simulation study, we conclude that DGLM is preferred over the BFT for practical use in plant vGWAS, although measures should be taken to overcome the severe loss in true positive detection rates when vQTNs have low MAFs. Future simulation studies should focus on more plant species, other quantitative genetics parameters, and additional statistical models. This could ultimately result in a more definitive understanding of the extent to which vGWAS could illuminate other genetic phenomena, including epistasis and GxE. To facilitate such future exploration of vGWAS approaches, the computational approaches we used to simulate traits controlled by vQTN are now publicly available free of charge in the simplePHENOTYPES R package (Fernandes and Lipka, 2020).

## Data Achieving

The genotypic data, simulated trait data, and code to simulate traits are available at https://github.com/mdm10-code/vGWAS_arabidopis_maize.

## Acknowledgements

We would like to thank reviewer one, reviewer two, and other reviewers for their helpful insights on vQTLs. We would also like to thank Dr. Hui Li for providing a rough Vv estimates for our oil-like settings. The research conducted in this manuscript is supported by the National Science Foundation project accession number 1355406, the University of Illinois Urbana-Champaign Department of Crop Science’s J.C. Hackleman and Lawrence E. Schrader and Elfriede Massier Plant Physiology Fellowship Programs.

